# Predicting the functional effects of voltage-gated potassium channel missense variants with multi-task learning

**DOI:** 10.1101/2021.12.02.470894

**Authors:** Christian Malte Boßelmann, Ulrike B.S. Hedrich, Peter Müller, Lukas Sonnenberg, Shridhar Parthasarathy, Ingo Helbig, Holger Lerche, Nico Pfeifer

## Abstract

**Purpose:** Variants in genes encoding voltage-gated potassium channels are associated with a broad spectrum of neurological diseases including epilepsy, ataxia, and intellectual disability. Knowledge of the resulting functional changes, characterized as overall ion channel gain- or loss-of-function, is essential to guide clinical management including precision medicine therapies. However, for an increasing number of variants, little to no experimental data is available. New tools are needed to evaluate variant functional effects.

**Methods:** We catalogued a comprehensive dataset of 959 functional experiments across 19 voltage-gated potassium channels, leveraging data from 782 unique disease-associated and synthetic variants. We used these data to train a taxonomy-based multi-task learning support vector machine (MTL-SVM), and compared performance to a baseline of standard SVMs.

**Results:** MTL-SVM maintains channel family structure during model training, improving overall predictive performance (mean balanced accuracy 0.729 ± 0.029, AU-ROC 0.757 ± 0.039) over baseline (mean balanced accuracy 0.645 ± 0.041, AU-ROC 0.710 ± 0.074). We can obtain meaningful predictions even for channels with few known variants (*KCNC1, KCNQ5*).

**Conclusion:** Our model enables functional variant prediction for voltage-gated potassium channels. It may assist in tailoring current and future precision therapies for the increasing number of patients with ion channel disorders.

## Introductions

Genetic disorders caused by variants in genes encoding voltage-gated ion channels are associated with a broad and heterogeneous spectrum of hereditary neurological, cardiac, nephrological, and other diseases.^1^ Among the ion channels associated with human disease, voltage-gated potassium channels (K_V_ 1–12) constitute the largest and most diverse super-family with regards to their structure, gating kinetics, expression profiles, pharmacology, and associated phenotypes. Variants in these channels are involved in developmental and epileptic encephalopathy (*KCNA2, KCNB1, KCNQ2, KCNQ3*), benign familial neonatal seizures (*KCNQ2, KCNQ3*), episodic ataxia or paroxysmal dyskinesia (*KCNA1*), nonsyndromic hearing loss (*KCNQ1, KCNE1, KCNQ4*), and long or short QT syndrome (*KCNQ1*).^2-5^ Many of these disease associations have only very recently been described, such as *KCNQ5*-associated intellectual disability, epileptic encephalopathy, or genetic generalized epilepsy, as well as *KCNC1*-associated progressive myoclonus epilepsy, intellectual disability, isolated myoclonus, or ataxia.^6-8^

In channelopathies, genetic and clinical data are inherently intertwined with the causative functional evidence provided by electrophysiological experiments. Voltage-clamp or patch-clamp studies in heterologous or neuronal expression systems are currently the methods of choice to bridge the translational gap between genetic diagnostics and clinical presentation, by establishing variant pathogenicity and elucidating the mechanism of channel dysfunction at a molecular and cellular level. For example, a variant may alter expression level, membrane trafficking, or the gating properties of a variant channel. The functional outcome can, in most cases, be characterized as an overall gain-of-function (GOF) or loss-of-function (LOF) with respect to subunit and channel function, with potential downstream effects on neuronal firing rate or network (dis)inhibition.^9^ However, we have to consider that cases exist, in which a net gain- or loss-of-function can be difficult to determine and may even differ between the type of neurons investigated.^10,11^ For clear net GOF or LOF cases, representing the majority of disease-associated variants, functional studies contribute to variant classification and inform clinical management, including genetic counselling and precision therapies.^12^ These cases often show clear phenotype-genotype correlations, facilitating diagnosis and offering therapeutic and prognostic guidance. But perhaps most interestingly, recent advances aim to correct the underlying biophysical defects. For example, retigabine selectively enhances the M-current in *KCNQ2-5* by stabilizing the open-pore conformation, which reverses the effects of some LOF variants.^13,14^ Conversely, 4-aminopyridine acts as a non-selective blocker of K_V_1.x channels and may be a safe and effective treatment option in patients with developmental and epileptic encephalopathy due to GOF variants in *KCNA2*.^15^

However, the unprecedented pace of genetic discoveries has surpassed the ability of traditional methods to provide sufficient causal evidence for variant interpretation.^16^ Consequently, there is an increasing number of variants for which no direct (i.e. specific variant studied in the same channel), indirect, or no functional data at all are available. Electrophysiological studies present a bottleneck in translational research: they are laborious, expensive and time-intensive, and therefore limited to the analysis of just a few variants at a time. High-throughput automated electrophysiology platforms for large-scale functional studies are currently in development and may ameliorate some of these disadvantages.^17^ But this also presents an opportunity to develop predictive frameworks that integrate computational, biochemical, evolutionary, and experimental sources of information.

Here, we present a machine learning-based model that leverages data from electrophysiological experiments to predict the functional effects of non-synonymous missense variants in voltage-gated potassium channels. Our model efficiently integrates data from two very different sources – synthetic variants from biophysical studies designed to understand channel function, as well as patient variants from clinical genetic testing. We demonstrate that taxonomy-based multi-task learning enables us to preserve information on the relatedness of ion channels, improving predictive performance. Our framework for functional variant prediction is scalable and can easily be extended to other use cases.

## Methods

### Data

Our dataset consisted of previously published, publicly available data from patch-clamp or voltage-clamp experiments, with each experiment describing the functional effects of a non-synonymous missense variant in a voltage-gated potassium channel. Many of these variants were previously found to be disease-causing in patients and hence functionally characterized. Notably, we also included data from non-human channels (Shaker, *KCNAS*) and from experiments that studied synthetic variants to understand biophysical channel function and structure-function relationships. Although these variants may not appear in disease context or in the general population, they increase the sample size and contain salient information which is useful for our model to learn. For each experiment, we collected the biophysical properties of the mutant channels, including information on the gating kinetics, peak current or conductance, and expression level or trafficking defects, if available. We used this information to assign one label to each variant, the label categorizing the net overall effect on ion channel function at subunit level: gain-of-function (GOF), loss-of-function (LOF), and no functional effect (neutral).

In total, we curated the results of 959 electrophysiological experiments from 163 publications. Of these, 22 experiments were excluded due to conflicts if contradicting functional effects for the same variant were described in two or more independent publications. For 67 experiments, we were unable to ascertain a label due to mixed or unclear functional effects without clear net gain- or loss-of-function. These variants, too, were excluded from the training set. Finally, 88 experiments were marked as duplicate and removed from the data set if two or more experiments reported the same functional effect for the same variant. The final dataset used for model training included 782 unique non-synonymous missense variants across 19 voltage-gated potassium channels: 165 GOF, 544 LOF and 73 neutral (Supplementary Figure 1). The dataset was independently validated by two expert electrophysiologists, and disagreements were resolved by consensus-based discussion.

### Features

Each variant was identified by its channel, sequence position, and amino acid substitution. From there, we extracted sequence- and structure-based features as follows: The residue-level properties of the original and substituted amino acids were included as features by physiochemical encoding, representing them in a lower-dimensional property space.^18^ Hydrophobicity was included as a separate feature, by principal component analysis of 98 hydrophobicity scales from Simm et al.^19^ Frequency and radicality of amino acid substitutions were described by the BLOSUM62 substitution matrix and Grantham score.^20,21^ Paralog conservation scores were obtained from PER viewer.^22^ Predictions of structural features were computed with PredictProtein, NetSurfP-2.0, and IUPred2A.^23-25^ This included information on secondary structure, residue accessibility, conservation, disordered regions, and interaction sites. We then annotated domains and motifs through UniProt and expert knowledge, including rich information on the position of gating charges, pore helix, S4-S5 linker, selectivity filter, PVP motif, and others.^26^ For preprocessing of our data, we used one-hot encoding of categorical features and scaling of numerical features.

### Models

In the previous steps, we have obtained a data matrix X with n observations (variants) by p dimensions (features). Each observation n belonged to one of three classes *y*_1_, …, *y*_*n*_ ∈ {*LOF,Neutral, GOF*} and to one of 19 tasks (voltage-gated potassium channels) *t*_1_, …, *t*_*n*_ ∈ {*KCNA*1, *KCNA*2, …, *KCNQ5*}. For a previously unseen variant x ∈ X we wanted to predict the functional effect, class label *ŷ*. Intuitively, if a new variant is similar to a known variant, it should receive the same class label. As classifier for this multiclass problem, we chose a support vector machine (C-SVM) with a radial basis function (RBF) kernel *K*_*n*_ with 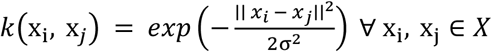. We took measures to reduce overfitting with *k*-fold cross validation, *k* = 10, and the cost regularization hyperparameter *C*. To correct for class imbalance, we assigned class weights *w* inversely proportional to the frequency of class *t, w*_*t*_ = *n*_*all*_/*n*_*t*_. The minority class is assigned a larger *w*_*t*_ and smaller C, thus increasing the misclassification penalty. The optimal values for the hyperparameters were estimated via grid search, with σ = {1e − 5, 1e − 4, …, 1e0} and C = {1e − 4, 1e − 2, …, 1e4}. We used Platt scaling to transform the output of our non-probabilistic binary linear classifier into a distribution of class probabilities, extended for the multiclass case. The model performance estimates, i.e. the expected generalization performance of the model on previously unseen data, were obtained by k-fold cross validation – a technique in which the model is trained and tested on different random partitions of the data throughout multiple exhaustive iterations, with the aim to obtain performance estimates that are more accurate and stable.

We constructed two methods for baseline comparison. For the single method, one SVM was trained for all observations belonging to one task, which yields *t* task-specific models. Each model was independently tuned and assessed. We did not attempt to train a model if there were less than 20 observations for the task, or if all observations had been assigned the same class label (e.g., channels for which only loss-of-function variants are known). In both cases, the resulting models would not have been meaningful. For the base method, one SVM was trained on all observations, irrespective of task membership. To compare our methods, we report the following metrics: i) Balanced Accuracy (BA), defined as the average of sensitivity and specificity for each class; ii) Matthews correlation coefficient (MCC), a symmetric correlation coefficient between true and predicted classes; and iii) Cohen’s Kappa, a measure of agreement between classifiers. The multi-class extension of these metrics has been described elsewhere.^28^ The area under the receiver operator characteristics (AU-ROC) curve was extended to the multi-class case using the method described by Hand and Till.^29^ We calculated the area under the precision-recall curve (AU-PRC) by taking the average of a one-versus-all approach. Further model information is available in our AIMe Registry report (https://aime.report/IFtQVF).

### Multi-task learning

To define our hierarchy of tasks *T*, we first created a multiple sequence alignment of all tasks with MUSCLE.^30^ For each observation, we noted the relative position on the consensus sequence and included this as an additional feature for all models. Each pairwise distance on the multiple sequence alignment *d*_*s,t*_ was transformed to a pairwise similarity measure by *γ*_*s,t*_ = *a* − *d*_*s,t*_/*d*_*max*_ to obtain our task similarity matrix *K*_*t*_. Here, *a* was a hyperparameter controlling the baseline similarity between tasks and was included in our grid search with *a* = {1, 3, 5, 10, 100}, while d_max_ was the maximum distance between two tasks in hierarchy *T*. This approach to taxonomy-based multitask learning is further detailed in Widmer et al. and is motivated by the previous work of Jacob and Vert.^31,32^ The final multi-task learning kernel matrix *K*_*m*_ was obtained by element-wise multiplication of the task kernel matrix *K*_*t*_ with our RBF kernel matrix *K*_*n*_. Intuitively, *K*_*m*_ embeds information on task similarity and instance similarity into the kernel matrix. The task hierarchy *T* and kernel matrix *K*_*t*_ for voltage-gated potassium channels are shown in Supplementary Figure 1.

### Software

This study was carried out in the R programming language, version 4.1.0, with RStudio, version 1.4.1106. Packages used include tidyverse (data preparation, preprocessing and visualization), the Bioconductor collection (feature extraction and multiple sequence alignment), kernlab and the e1071 interface for LIBSVM (model training), yardstick and pROC (model assessment), and Shiny (graphical user interface and web application). R Session Information is included as part of our data availability statement.

## Results

For our first naïve baseline method, we trained one separate SVM for variants of each voltage-gated potassium channel (single method). For tasks with sufficiently large and balanced training data sets, these single channel-specific models were already feasible, such as for variants in *KCNA2* (BA 0.689, MCC 0.293), *KCNQ1* (BA 0.762, MCC 0.321), or *KCNQ4* (BA 0.813, MCC 0.489). However, we did not obtain meaningful models for 6/19 channels (Figure 1). For these channels, either an insufficient sample size of variants was available for training, or each variant had been assigned the same label (e.g. only loss-of-function variants). Notably, this included channels that have only recently been associated with human disease, such as *KCNC1* or *KCNQ5*, for which little functional data are available. The single method is thus limited to a few common channels, but performs reasonably well on average (Table 1).

**Figure 1.**
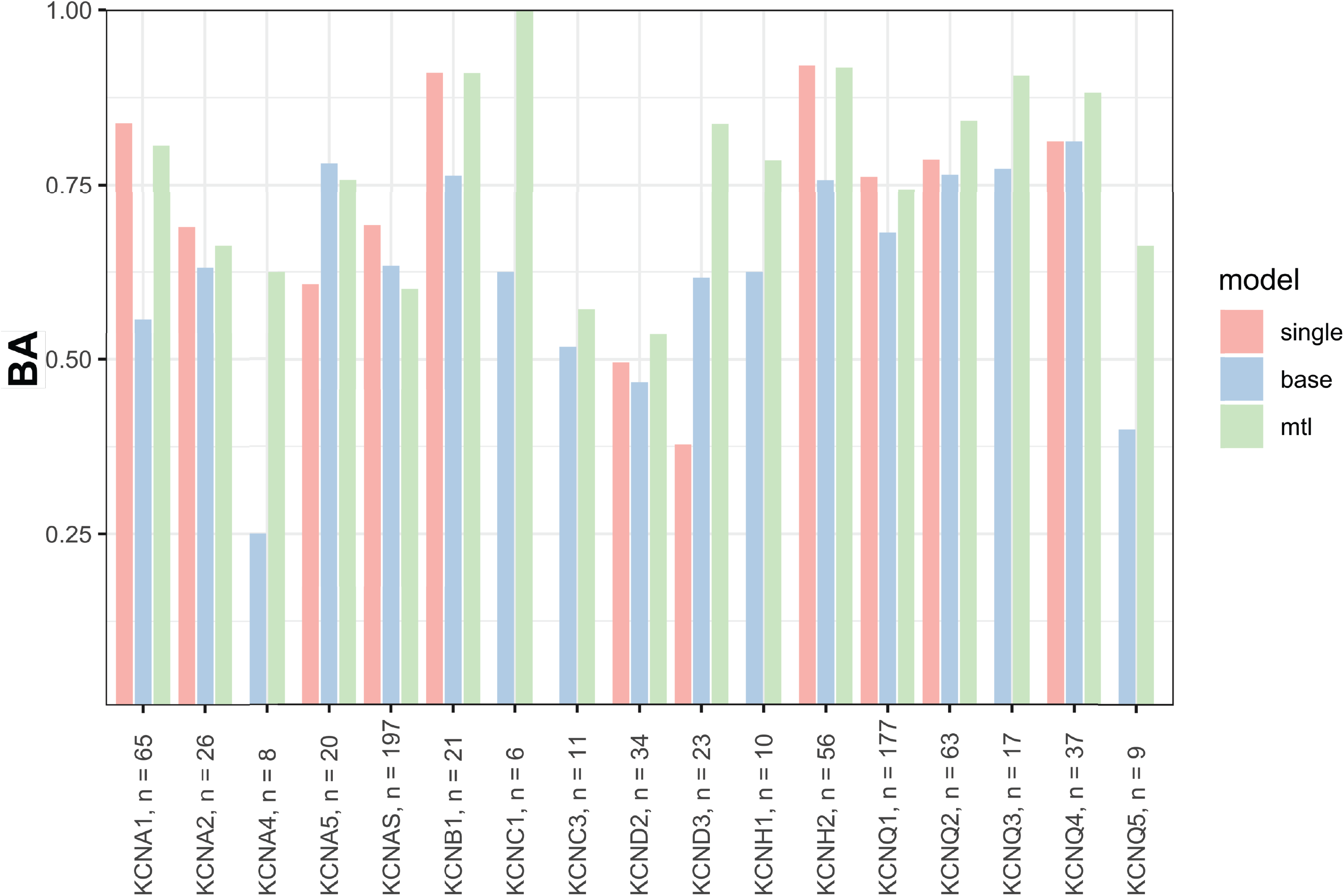
Task-specific cross-validation model performance estimate of mean Balanced Accuracy for each method. Channels are given with their respective number of variants (n) included in the training data set. Two tasks, *KCNC2* and *KCNH5*, contained just one variant each and are not included in the performance estimate. Abbreviations: BA – Balanced Accuracy.

**Table 1.** Cross-validation model performance estimates for each method. Metrics are reported as mean ± standard deviation, where applicable. Metrics are calculated either over all variants or over all tasks. Tasks are further divided in subgroups of common (*KCNA1, KCNA2, KCNA5, KCNAS, KCNB1, KCND2, KCND3, KCNH2, KCNQ1, KCNQ2, KCNQ4*) and rare (*KCNA4, KCNC1, KCNC2, KCNC3, KCNH1, KCNQ3, KCNQ5*) channels, depending on variant frequency in the training data set (cut-off ≥20 variants). For these subgroups, BA and AU-ROC are shown. Abbreviations: AU-PRC – area under the precision-recall curve; AU-ROC – area under the receiver operator characteristic; BA – Balanced Accuracy; MCC – Matthews correlation coefficient.

|  | single |  |  |  |  | base |  |  |  |  | mtl |  |  |  |  |
| --- | --- | --- | --- | --- | --- | --- | --- | --- | --- | --- | --- | --- | --- | --- | --- |
| | BA | Cohen's $\kappa$ | MCC | AU-ROC | AU-PRC | BA | Cohen's $\kappa$ | MCC | AU-ROC | AU-PRC | BA | Cohen's $\kappa$ | MCC | AU-ROC | AU-PRC |
| All variants | 0.742 ± 0.023 | 0.295 ± 0.056 | 0.297 ± 0.057 | 0.734 ± 0.031 | 0.513 ± 0.041 | 0.645 ± 0.041 | 0.205 ± 0.079 | 0.225 ± 0.086 | 0.710 ± 0.074 | 0.501 ± 0.062 | 0.729 ± 0.029 | 0.331 ± 0.054 | 0.345 ± 0.055 | 0.757 ± 0.039 | 0.553 ± 0.041 |
| All tasks | 0.719 ± 0.163 | 0.234 ± 0.117 | 0.278 ± 0.149 | 0.529 ± 0.186 | 0.488 ± 0.167 | 0.627 ± 0.153 | 0.191 ± 0.139 | 0.323 ± 0.185 | 0.674 ± 0.183 | 0.500 ± 0.209 | 0.767 ± 0.138 | 0.238 ± 0.189 | 0.385 ± 0.241 | 0.743 ± 0.139 | 0.530 ± 0.264 |
| Common tasks | 0.719 | – | – | 0.539 | – | 0.643 | – | – | 0.675 | – | 0.760 | – | – | 0.735 | – |
| Rare tasks | – | – | – | – | – | 0.588 | – | – | 0.667 | – | 0.785 | – | – | 0.833 | – |

Next, we trained one SVM on a pooled dataset of all variants across the voltage-gated potassium channel family (base method). This represents a conventional single-task approach, for which data is shared between all tasks without differentiation. Clearly, sharing data is beneficial, enabling us to obtain predictions for variants from all 19 channels in our dataset. However, predictive performance was low for some channels, e.g. *KCNA4, KCND2*, and *KCNQ5* (Figure 1). We also observed that the task-specific model performance estimate of the base method was lower than that of some single SVMs, and deteriorated further for some rare channels (Table 1). Put in other words, adding data from other channels generally caused the model to perform worse than a single, channel-specific model. It stands to reason that one global model cannot fit all these channels equally well, some of which are markedly different from each other in structure and function. The average overall performance of this base method was not sufficient to be clinically useful (Table 1).

Our multi-task learning approach (mtl method) embeds structural and evolutionary pairwise similarity between ion channels in the kernel matrix, efficiently integrating prior knowledge across all channels and channel sub-families. Intuitively, the model learns to assign more weight to variants from the same or similar channels, and less or no weight to variants from more distantly related channels. At the same time, transfer learning of a shared representation allows the model to learn higher-level concepts that apply across all channels, such as the relative feature importance of substitution radicality or residue conservation. Thus, this method allowed us to obtain meaningful predictions for variants from every channel, while maintaining a good overall performance (Figure 1). More importantly, multi-task learning achieved strong performance even for variants in rare channels (BA 0.785 and AU-ROC 0.833 versus BA 0.588 and AU-ROC 0.667 for the base method, Table 1). The superior performance of the multi-task learning method over the base method is also shown as an overall gain in AU-ROC (mean AU-ROC 0.757 ± 0.039 versus AU-ROC 0.710 ± 0.074) (Figure 2).

**Figure 2.**
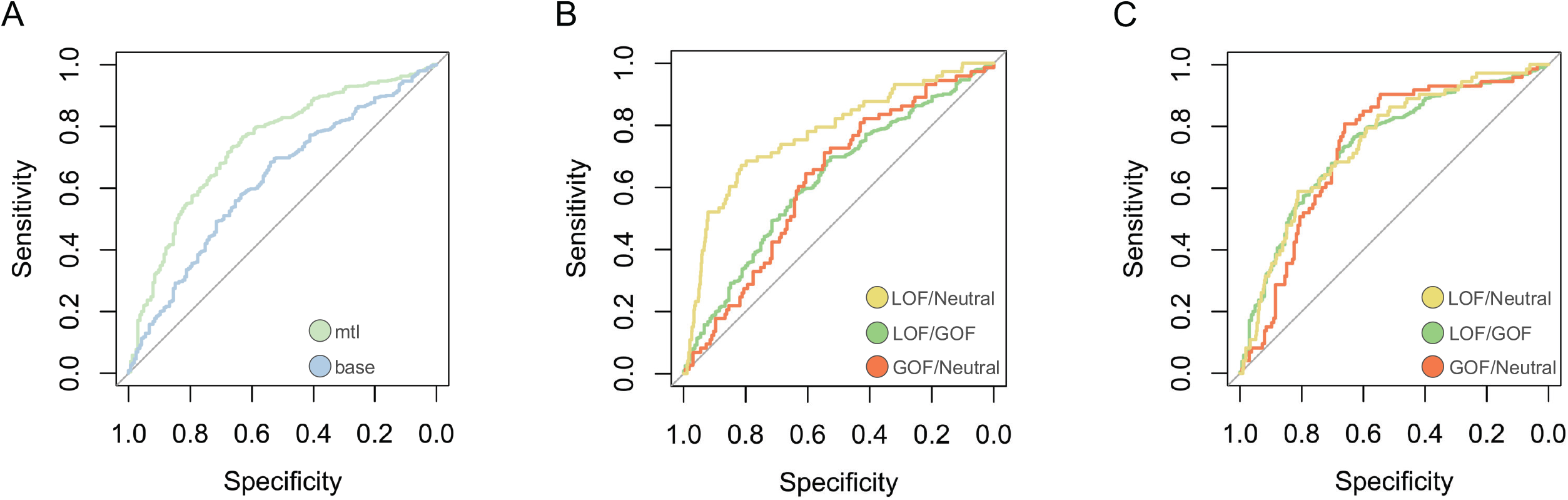
ROC of the baseline (“base”) versus multi-task learning (“mtl”) models, AU-ROC 0.710 ± 0.074 versus AU-ROC 0.757 ± 0.039. *A*: The multi-task learning model better discriminates between GOF and LOF variants. *B:* ROC curves for the baseline model. *C*: ROC curves for the multi-task learning model. Abbreviations: AU-ROC – area under the receiver operating characteristic; GOF – gain-of-function; LOF – loss-of-function.

Additionally, some new missense variants in voltage-gated potassium channels have very recently been described, which have not yet been included in the training set. Firstly, Miceli et al. described four heterozygous de novo variants in *KCNA1:* P403S, P405L, P405S and A261T.^33^ Our MTL-SVM correctly predicted three out of four variants with high confidence (Figure 3A). The A261T variant, which resulted in a hyperpolarizing shift of the voltage-dependence of activation (GOF), was incorrectly predicted as LOF. This is expected behavior: Gain-of-function in this channel, previously only seen in some synthetic variants from biophysical research, is very rare and has only now been associated with disease-causing variants. Clearly, experimental evidence still remains crucial to generate causative evidence. Secondly, Imbrici et al. described a heterozygous de novo variant in *KCNA2*, E236K.^34^ This variant caused a hyperpolarizing shift of the voltage-dependence of activation and slowed kinetics of activation and deactivation, resulting in an overall mixed GOF/LOF. Variants without a clear net overall functional effect were excluded from our model’s training set. Still, the model correctly recognized that E236K is equally likely to result in a GOF or LOF (Figure 3). For both use cases presented here (*KCNA1, KCNA2*), possible precision therapies based on variant function have been proposed.^15,33,34^

**Figure 3.**
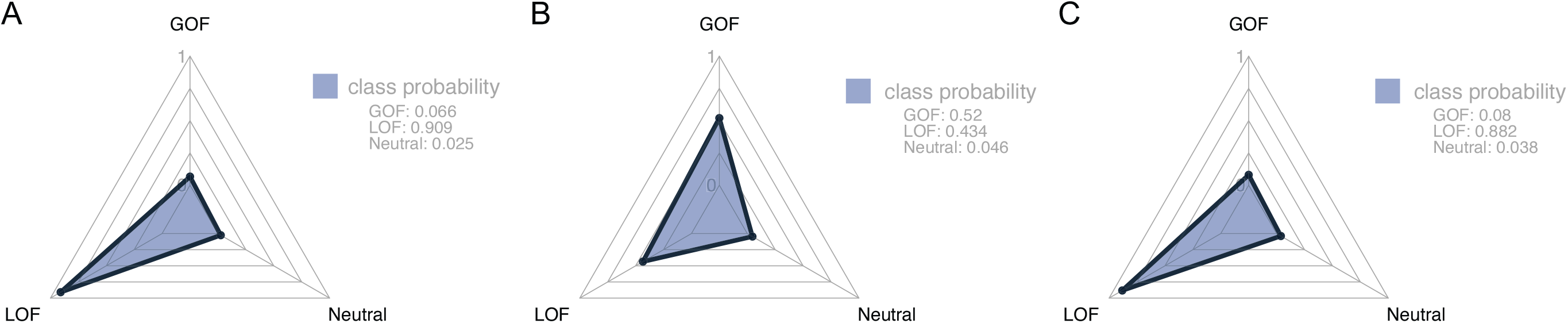
Radar plots of class probability distributions, with class probabilities shown in the legend. A: *KCNA1* P403S, P405L and P405S were correctly predicted as LOF (P405L and P405S are not shown). B: For *KCNA2* E236K, a variant with mixed functional effects, the model correctly predicted an equal probability of GOF and LOF. C: *KCNA1* A261T was incorrectly predicted as LOF.

## Discussion

Next-generation sequencing continues to generate large datasets of genetic variants at an ever-increasing pace, driven by both clinical genetic diagnostics and the international cooperation of sequencing consortia. *In silico* tools can aid in predicting the pathogenicity of missense variants of unknown significance. Some of these prediction models are tailored to specific genes, such as Q1VarPred for *KCNQ1*, which outperforms other methods for pathogenicity prediction by incorporating some functional data.^35^ A gene-family-specific tool, KvSNP, outperforms other methods for pathogenicity prediction across all voltage-gated potassium channels.^36^ However, none of these tools are designed to provide information on the alteration of biophysical function at a molecular and cellular level: This is the difference between traditional pathogenicity prediction and functional variant prediction. In ion channel disorders, this functional knowledge is crucial to guide diagnosis, treatment, and prognosis.

Few studies have attempted to address this unmet need. Recently, Heyne et al. presented funNCion, a statistical method for functional variant prediction of missense variants in voltage-gated sodium (Na_V_) and calcium channels (Ca_V_).^37^ In these channel families, *de novo* variants are enriched at paralog-conserved sites. Heyne et al. hence pooled variants for further analysis, under the assumption of similar molecular mechanisms across individual channels. Their training data set included 827 LOF and GOF variants in Na_V_ and Ca_V_, where the functional label was inferred from phenotype data alone (e.g., age at seizure onset). Using a Gradient Boosting machine, they predicted an independent test set of 87 variants with an estimated performance of BA 0.73, AU-ROC 0.73, and MCC 0.45. Interestingly, they reported a similar performance when predicting Na_V_ variants with a training set of either Na_V_ alone, or both Na_V_ and Ca_V_. Conversely, the prediction on a test set of Ca_V_ was worse when the training set contained just Ca_V_ alone, while performance improved when training on just Na_V_, or both Na_V_ and Ca_V._ They concluded that the increased power obtained by combining Na_V_s and Ca_V_s outweighs the differences between these channel types. Evidently, pooling data may be beneficial. But how do we formalize the general notion of difference between ion channels, and how do we then decide to which variants in the training data we want to assign more weight?

The answer lies in the notion of an implicit or latent task relatedness. If we consider each member of the family of voltage-gated potassium channels as a single task, we can then employ multi-task learning, in which information is shared between the tasks: We learn our tasks in parallel using a shared representation, which has been shown to outperform the conventional single-task learning methods introduced above.^38^ Task relatedness may be learnt from the data structure, but we may also view this as an opportunity to introduce prior domain knowledge. For example, Jacob and Vert used a user-defined or supertype-based measure of similarity to improve peptide-MHC-I binding models across different alleles.^32^ They demonstrated that the method can be decomposed into two steps: i) Choosing an explicit description or pairwise similarity kernel for our tasks (channels) and observations (variants); ii) Applying an SVM to the product kernel. For our task kernel, we chose the taxonomy-based method by Widmer et al., for which task relatedness is derived from the hierarchical structure of a phylogeny tree, which is readily obtained through multiple sequence alignment.^31^ This is the same principle hierarchy that motivates the classification of voltage-gated potassium channel families, and it appropriately separates channels both according to their function (i.e. different potassium currents) and their tissue expression levels.^39^

Here, we have demonstrated the feasibility of functional variant prediction in voltage-gated potassium channels, supported by a considerable experimental data set. We then introduced an MTL-SVM framework which is uniquely suited to handle predictions across large and diverse channel families. Our multi-task learning method improves upon two single-task learning baseline models and compares favorably to the method by Heyne et al. (2020). Notably, we achieved good predictive performance even in channels for which data from functional studies are sparse, such as *KCNC1* or *KCNQ5*, which have only recently been associated with human disease.^6,7^ Thus, our method may prospectively enable predictions for emerging channels and new disease-associated variants as they become relevant. Another strength of our approach is that we can leverage data from non-human channels and synthetic variants, making efficient use of prior information on structure-function relationships from decades of biophysical research to improve model performance. This tool may generate supportive evidence for in silico variant interpretation, and may assist clinicians and researchers in generating new hypotheses about voltage-gated potassium channel disorders, including potential new applications for precision therapies.

Importantly, our model predicts functional changes on a main/alpha subunit level and does not include information on modifying subunits (*KCNFx, KCNGx, KCNVx, KCNSx*, etc.) or experimental metadata (e.g., expression system, recording protocol). Care should be taken when generalizing the results of the prediction to a neuronal, network, or phenotype level. In silico predictions should always be considered in concert with other available sources of evidence. By themselves, in silico predictions are not considered strong or causal evidence by the genetic testing standards established by the American College of Medical Genetics and Genomics.^40^ We have assembled a large and representative data set of variants across the superfamily of voltage-gated potassium channels but, given their diversity and the wealth of published literature, this data set is far from complete. There is some concern for selection bias, as electrophysiological experiments are more likely to study either pathogenic variants, or those in locations of structural and functional importance. Further work is clearly required to improve model performance. These methods will benefit from large homogeneous data sets provided by high-throughput electrophysiology, as well as standardized experimental and phenotype data curation and integration frameworks that are currently in development. Additionally, static and dynamic structural information from AlphaFold and molecular dynamics may prove useful.

## Data Availability

Data and code generated during this study, including pre-processed training data and pre-trained models, are available on Zenodo (DOI 10.5281/zenodo.5749260).

## Acknowledgements

This work was supported by intramural funding of the Medical Faculty, University of Tuebingen (PATE F.1315137.1), the Federal Ministry for Education and Research (Treat-ION, 01GM1907A/B/G/H) and the German Research Foundation (FOR-2715, Le1030/16-2, He8155/1-2). The funders had no role in study design, data collection, data analysis, and decision to prepare or publish the manuscript.

The authors would like to thank Mahmoud Koko (Department of Neurology and Epileptology, Hertie Institute for Clinical Brain Research, University of Tübingen, Tübingen, Germany) for his assistance in data preparation for *KCND2/KCND3* variants.

## Author Information

Resources: H.L., N.P.; Funding acquisition: C.M.B., H.L., N.P.; Supervision: H.L., N.P.; Methodology: C.M.B.; Data Curation: C.M.B., U.B.S., P.M.; Validation: U.B.S., P.M.; Software: C.M.B.; Formal analysis: C.M.B., L.S., S.P., I.H.; Writing – Original Draft: C.M.B.; Visualization: C.M.B.; Writing – Review & Editing: L.S., I.H., H.L., N.P.

## Ethics Declaration

Not applicable.

## Abbreviations

AU-PRC: area under the precision-recall curve
AU-ROC: area under the receiver operator characteristic
BA: Balanced Accuracy
GOF: gain of function
LOF: loss of function
MCC: Matthews correlation coefficient
MTL: multi-task learning
RBF: radial basis function
SVM: support vector machine

## Figure legends

**Supplementary Figure 1.**
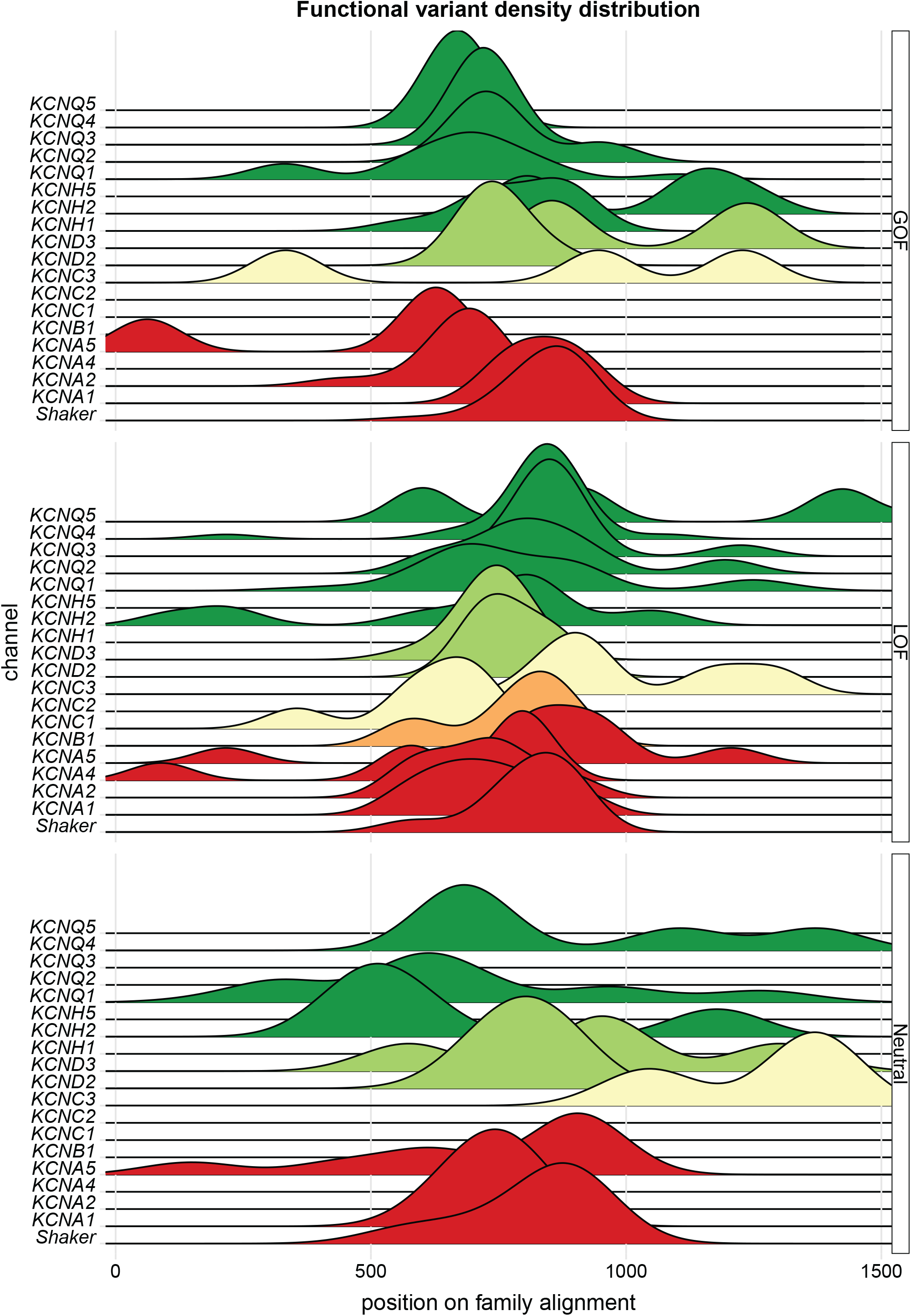
Functional variant density distribution along the channel super-family multiple sequence alignment. Variants are grouped by functional effect (GOF, LOF and Neutral) and colored by their channel sub-family.

**Supplementary Figure 2.**
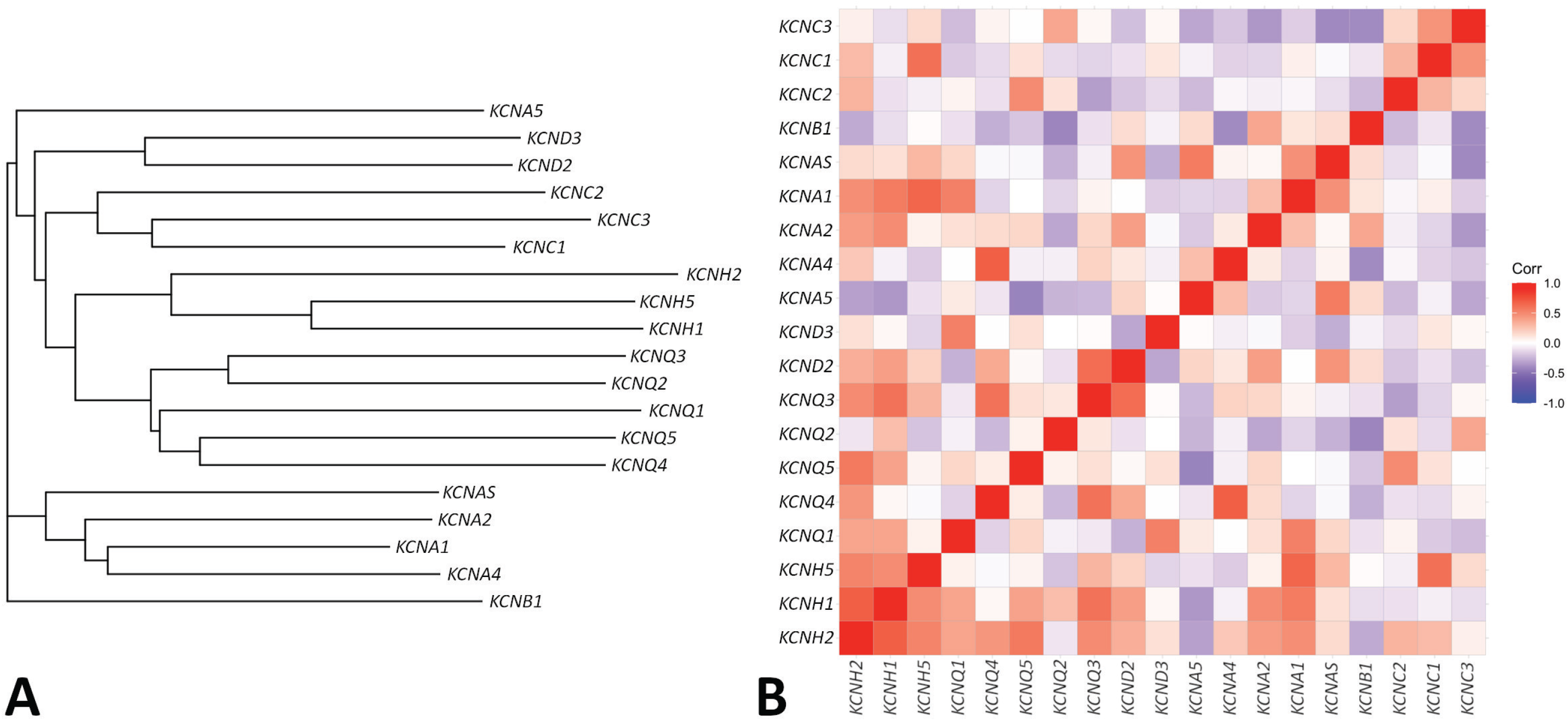
*A*: Task hierarchy *T* derived from MUSCLE multiple sequence alignment. *B*: Visualization of the task similarity matrix *K*_*t*_ as a correlation plot, where positive values (shown in red) correspond to a lower pairwise distance and higher pairwise similarity, and vice versa.

